# Calcium-Permeable AMPA Receptor Activity and GluA1 Trafficking in the Basolateral Amygdala Regulate the Positive Reinforcing Effects of Alcohol

**DOI:** 10.1101/2021.03.25.436994

**Authors:** Sara Faccidomo, Elizabeth S Cogan, Olivia J. Hon, Jessica L Hoffman, Briana L Saunders, Vallari R Eastman, Michelle Kim, Seth M. Taylor, Zoé A McElligott, Clyde W Hodge

**Affiliations:** Bowles Center for Alcohol Studies; Neuroscience Curriculum; Department of Psychiatry; Department of Pharmacology, The University of North Carolina at Chapel Hill, Chapel Hill, NC 27599

## Abstract

Addiction is viewed as maladaptive glutamate-mediated neuroplasticity that is regulated, in part, by calcium-permeable AMPA receptor (CP-AMPAR) activity. However, the contribution of CP-AMPARs to alcohol-seeking behavior remains to be elucidated. We evaluated CP-AMPAR activity in the basolateral amygdala (BLA) as a potential target of alcohol that also regulates alcohol self-administration in C57BL/6J mice. Operant self-administration of sweetened alcohol increased spontaneous EPSC frequency in BLA neurons that project to the nucleus accumbens as compared to behavior-matched sucrose controls indicating an alcohol-specific upregulation of synaptic activity. Bath application of the CP-AMPAR antagonist NASPM decreased evoked EPSC amplitude only in alcohol self-administering mice indicating alcohol-induced synaptic insertion of CP-AMPARs in BLA projection neurons. Moreover, NASPM infusion in the BLA dose-dependently decreased the rate of operant alcohol self-administration providing direct evidence for CP-AMPAR regulation of alcohol reinforcement. Since most CP-AMPARs are GluA1-containing, we asked if alcohol alters the activation state of GluA1-containing AMPARs. Immunocytochemistry results showed elevated GluA1-S831 phosphorylation in the BLA of alcohol as compared to sucrose mice. To investigate mechanistic regulation of alcohol self-administration by GluA1-containing AMPARs, we evaluated the necessity of GluA1 trafficking using a TET-ON AAV encoding a dominant-negative GluA1 c-terminus (GluA1ct) that blocks activity-dependent synaptic delivery of native GluA1-containing AMPARs. GluA1ct expression in the BLA reduced alcohol self-administration with no effect on sucrose controls. These results show that CP-AMPAR activity and GluA1 trafficking in the BLA mechanistically regulate the reinforcing effects of sweetened alcohol. Pharmacotherapeutic targeting these mechanisms of maladaptive neuroplasticity may aid medical management of alcohol use disorder.

## 1 INTRODUCTION

Alcohol use disorder (AUD) is one of the most widespread neuropsychiatric conditions with a lifetime prevalence of 29.1% (men 36%; women 22.7%) in the United States ^1^. Although it is widely recognized that chronic repetitive alcohol use is fundamental to AUD, the neural and behavioral mechanisms that contribute to this pathological process remain to be fully elucidated. The development and progression of AUD is engendered by pathological repetitive alcohol use. Derived from Thorndike’s *Law of Effect* ^2^, the fundamental behavioral process of *reinforcement* reflects the tendency of all animals, human and non-human, to repeat responses that produce a desired outcome ^3^. Thus, evidence indicates that positive reinforcing effects of alcohol drive repetitive drug use during the initial binge-intoxication stage of addiction and promote the development of dependence ^4,5^. For this reason, research that elucidates the neural mechanisms of the positive reinforcing properties of alcohol has potential to move the field forward in understanding the etiology and progression of AUD, and inform the development of pharmacotherapeutic strategies ^6^.

An abundance of preclinical evidence indicates that alcohol and other drugs of abuse gain long-lasting control over behavior, in part, by hijacking glutamatergic α-amino-3-hydroxy-5-methyl-4-isooxazole receptor (AMPAR) mechanisms of plasticity in key structures of the brain’s reward system, including the amygdala ^4,7-11^. Accordingly, systemic pharmacological activation of AMPAR activity increases the positive reinforcing effects of alcohol and promotes cue-induced reinstatement of alcohol-seeking behavior ^12^. Within reward circuits, activation of AMPARs in the amygdala promotes escalated operant alcohol self-administration in a CaMKII-dependent manner ^13^. Complementary loss of function evidence shows that inhibition of amygdala CaMKII or AMPAR activity specifically inhibited the positive reinforcing properties of alcohol but not sucrose ^14^. Since AMPAR-mediated plasticity in the amygdala is required for the development of new behavior (e.g., learning) and retention of actions (e.g., memory), these neural processes may also underlie the development and maintenance of chronic repetitive alcohol self-administration, which is a hallmark behavioral pathology that underlies AUD.

AMPARs are glutamate-gated ion channels comprised of combinations of GluA1 – 4 subunits. Their activity is determined by a variety of post-translational modifications including phosphorylation, trafficking, and subunit composition ^15^. Most glutamatergic synapses are populated by GluA2-containing, Ca^2+^-impermeable AMPARs (CI-AMPARs) made up from GluA1/GluA2 and GluA2/GluA3 heteromers ^16^. By contrast, certain plasticity inducing events ^17^, including exposure to drugs of abuse ^18-21^, can promote an activity-dependent increase in synaptic expression of CP-AMPARs. In some instances, this is associated with GluA1 phosphorylation at S831 and S845 ^22,23^. Our prior work has shown that alcohol self-administration is associated with upregulated AMPAR synaptic activity and increased pGluA1-S831 in the amygdala ^14,24^ suggesting that repetitive alcohol use may target CP-AMPAR activity in this brain region. This is consistent with evidence that chronic intermittent ethanol (CIE) exposure increases CP-AMPAR synaptic activity in the nucleus accumbens (NAcb) ^25,26^. However, it is not known if CP-AMPAR activity in the amygdala is a target of alcohol that mechanistically regulates self-administration behavior.

In this study, we evaluated the role of CP-AMPARs in the basolateral amygdala (BLA) in the positive reinforcing effects of sweetened alcohol. Sweetened solutions have been used in preclinical alcohol research for many years and offer a variety of advantages over unsweetened alcohol including enhanced face validity to the human condition where sweeteners are commonly added to alcohol during the initial binge-intoxication stage of addiction (reviewed by ^27^). Here, we utilized a well-characterized method that compares operant self-administration of sweetened alcohol (ethanol 9% v/v + sucrose 2% w/v) to parallel behavior-matched sucrose-only (sucrose 2% w/v) controls ^14,28-31^. Availability of a non-drug behavior-matched control is important when evaluating the role of CP-AMPARs in alcohol self-administration because the BLA sends numerous glutamatergic projections to reward-related brain regions including the NAcb ^32^ and response contingent optical activation of BLA-to-NAcb excitatory projection fibers reinforces nose-poking behavior ^33^, indicating that the BLA is an integral node in non-drug reward circuitry. Thus, by making direct comparisons to a non-drug reward that generates the same level of self-administration behavior, the present study has potential to shed light on the hypothesis that CP-AMPARs in the BLA are a target of alcohol that, in turn, regulate alcohol-seeking behavior via misappropriation of glutamatergic cellular mechanisms ^34,35^.

To address this goal, we first evaluated synaptic properties and response to NASPM as an index of CP-AMPAR accumulation in BLA-to-NAcb projection neurons using a retrobead strategy in ex vivo slices from sweetened alcohol or sucrose self-administering mice. To obtain direct evidence for CP-AMPAR regulation of alcohol self-administration, a site-specific pharmacological approach was taken to determine CP-AMPAR activity in the BLA is required for the positive reinforcing effects of alcohol as measured by operant self-administration. To determine if CP-AMPAR activity is a neural target of alcohol, we then assessed the impact of alcohol self-administration on pGluA1-S831 immunoreactivity in the BLA of C57BL/6J mice. Finally, since postsynaptic CP-AMPAR activity is determined, in part, by membrane trafficking of GluA1-containing AMPARs, we utilized a viral vector strategy to determine if GluA1 trafficking is required for the positive reinforcing effects of alcohol. A TET-ON adeno-associated viral vector (AAV) was expressed in the BLA that encodes a dominant-negative form of the AMPAR GluA1 c-terminus (GluA1ct), which has been shown to prevent activity-dependent synaptic delivery (membrane trafficking) of GluA1-containing AMPARs by blocking membrane insertion of the c-terminus of native GluA1-containing AMPARs ^36,37^. Together, these studies provide compelling evidence at the molecular, physiological, and behavioral level that the reinforcing effects of alcohol require CP-AMPAR activity and GluA1 trafficking in the BLA.

## 2 MATERIALS AND METHODS

Full details of experimental methods are in ***Supplemental Material***.

### 2.1 Mice

Adult male C57BL/6J mice (n=56) were group-housed (n=4/cage). All experimental groups were trained in cohorts of 8 mice. All experiments were approved by the Institutional Animal Care and Use Committee of the University of North Carolina at Chapel Hill and animals were cared for in accordance with the Guide for the Care and Use of Laboratory Animals ^38^.

### 2.2 Operant Self-administration Training and Baseline

Self-administration sessions were conducted in computer controlled two-lever operant conditioning chambers (Med Associates, St. Albans, VT) as previously reported ^14,28-31,39,40^. Briefly, lever-press responses were trained with sucrose-only (2% w/v; parallel non-drug control) or sweetened alcohol (9% v/v ethanol + 2% w/v sucrose) reinforcement for the duration of experiments. Reinforcement was maintained on a fixed-ratio 4 (FR4, 4 lever presses per reinforcer) schedule of reinforcement. After 30-40 daily 1-hr baseline sessions, separate cohorts of mice were used for 1) measurement of AMPAR GluA1-S831 phosphorylation in the BLA via immunohistochemistry; 2) whole-cell patch-clamp recordings of BLA-NAcb projection neurons; 3) site-specific infusion of the CP-AMPAR antagonist NASPM; or 4) expression of the dominant-negative GluA1ct AAV in the BLA to evaluate mechanistic regulation of self-administration. Mice that did not exhibit stable or sufficient levels of operant responding for either alcohol or sucrose (>10 reinforcements/hr) after surgery were excluded from the electrophysiology, microinjection and AAV studies.

### 2.3 Whole-cell patch-clamp recordings from NAcbC projecting BLA neurons

To examine the BLA to NAcb projecting neurons in alcohol vs. sucrose self-administering mice, fluorescent retrobeads (Lumafluor) were injected bilaterally in the NAcbC (0.3 µl/injection). Mice then self-administered alcohol (n=8 mice) or sucrose (n=8 mice) for 38-42-d as described above. Whole-cell voltage-clamp recordings were then conducted from retrobead-positive neurons in the BLA (24-h post alcohol or sucrose self-administration; NASPM: n=6 cells/6 mice, n=5 cells/5 mice, sEPSCs: n=6 cells/6 mice, n=6 cells/4 mice, respectively) as described previously ^41^ and detailed in *Supplemental Material*. Mouse attrition was due to failure to reestablish baseline behavior or presence of polysynaptic currents in the BLA that precluded analysis.

### 2.4 NASPM Microinjection in the BLA

To assess potential mechanistic regulation of alcohol self-administration by CP-AMPARs, the GluA2-lacking AMPAR antagonist NASPM (0, 1, 3, 5.6 or 10 μg / 0.5 µl) was prepared in aCSF vehicle and infused in the BLA immediately prior to a 1-h operant conditioning session in alcohol self-administering mice (n=6) as described ^14,30,42^. Each mouse received all doses of NASPM administered according to a within-subject balanced Latin Square design to control for dose order effects. Each subsequent injection only occurred when operant responding returned to baseline, and a maximum of 2 injections occurred each week, with a minimum of 48 hrs between each injection. No carryover effects of NASPM to post-injection days was observed. Behavioral measures including total lever-press responses and response rate were computer recorded. Bilateral injection sites were verified histologically to be in the BLA. To reduce the use of vertebrate animals, sucrose self-administration was not tested since NASPM had no effect on EPSC amplitude in the BLA of sucrose exposed mice. The repeated dosing design of this experiment is consistent with other studies evaluating the role of GluA2-lacking AMPARs in drug-associated reward using repeated site-specific infusion of NASPM ^19,43,44^.

### 2.5 Immunohistochemistry (IHC)

Coronal brain sections were prepared from alcohol (n=8) and behavior-matched sucrose (n=8) self-administering mice 24-h after the last session for quantification of pGluA1-S831 positive pixels/mm^2^ and cells/mm^2^ in the BLA using BIOQUANT image analysis software (BIOQUANT Image Analysis Corporation, Nashville, TN, USA). A representative coronal section through the BLA was co-labeled with GluA1 and Draq5 (labels nuclear DNA) to qualitatively evaluate the cytological expression of GluA1-containing AMPARs. All IHC procedures were conducted as previously described ^14,39,45-47^.

### 2.6 GluA1ct dominant-negative viral construct

The adeno-associated virus (AAV) construct used in the present study (pAAV-TRE-GluA1ctmCherry-CMV-Tet-On3G) was generated as previously described ^37^ kindly provided by Dr. Manuel Mameli. The GluA1ct AAV was fused to mCherry and inserted under the control of a TET-ON system ^37^. The AAV plasmids were made into recombinant AAV2/5 particles by the Vector Core Facility at the University of North Carolina, Chapel Hill, USA.

This well-characterized vector encodes a dominant-negative form of the GluA1 c-terminus (GluA1ct, corresponding to amino acids 809 to 889) that temporally prevents activity-dependent synaptic delivery of the AMPAR GluA1 subunit ^36^ via doxycycline-driven recombination. Electrophysiological properties of this vector have been characterized extensively and shown no effect on basal excitatory neurotransmission ^36,37^, but blockade of GluA1 synaptic delivery induced by LTP and inhibition of behavioral plasticity as indexed by fear conditioning following infection in the lateral amygdala ^36^. Inhibition of GluA1 trafficking via Tet-On expression of GluA1ct AAV in the lateral habenula prevented cocaine-evoked synaptic potentiation of AMPA currents ^37^.

### 2.7 GluA1ct AAV expression in the BLA

Mice were trained to self-administer alcohol (n=16) or sucrose-only (n=16) as a behavior-matched control. After 21 days of baseline performance, AAV-GluA1ct-mCherry (alcohol n=6; sucrose n=6) or mCherry control (alcohol n=8; sucrose n=4) were infused bilaterally in the BLA at a volume of 300nl/side (see *Stereotaxic Surgery in Supplemental Materials* for details). After 1 week of recovery, self-administration resumed for 15 sessions to assure a stable baseline and provide 3 weeks for validated GluA1ct expression ^37^. Mouse attrition was due to failure to recover baseline performance. Mice were then administered a palatable oral pellet containing doxycycline (0 or 10 mg/kg; 5 mg/kg twice per day at 0900 and 1800 hrs) for a total of 15 consecutive days. Consumption of the doxycycline pellet within 5-10 min was confirmed by experimenter observation. This twice per day doxycycline dosing regimen replicates the method of Meye et al (2015), which showed no impact of GluA1ct expression on basal excitatory synaptic activity (e.g., EPSC frequency, amplitude, or AMPA/NMDA ratio) but blockade of cocaine-induced GluA1 trafficking ^37^. Self-administration and locomotor behavior were monitored during the final 5 days of doxycycline exposure to evaluate potential mechanistic regulation by GluA1 trafficking in the BLA. After completion of the study, mice were deeply anesthetized, intracardially perfused, and brains were collected for histological verification.

### 2.8 Locomotor Activity

To address the possibility that changes in alcohol self-administration induced by NASPM infusion or GluA1ct AAV expression in the BLA were associated with nonspecific behavioral effects, spontaneous open-field locomotor activity (general motor function) and thigmotaxis (anxiety-like behavior) were assessed in separate 1-hr tests after completion of the self-administration dose-effect curves as reported previously ^31,48-52^.

### 2.9 Statistical Analysis

Data are presented as MEAN±SEM. All statistical analyses and graphic representations of data were performed using Prism software (version 9.0, GraphPad Software, San Diego, CA). Group comparisons were conducted via unpaired t-test, one-way RM ANOVA, or two-way RM ANOVA. Post hoc comparisons were conducted with Dunnett’s or Šídák’s multiple comparison procedures where appropriate. P values of < 0.05 were described as statistically significant.

## 3 RESULTS

### 3.1 Alcohol increases synaptic insertion of CP-AMPARs in BLA→NAcb projection neurons

To determine if operant alcohol self-administration is associated with postsynaptic accumulation of CP-AMPARs in projection neurons, whole-cell patch-clamp recordings were conducted from NAcb retrobead-positive BLA neurons 24-h after 40 days of operant alcohol vs. sucrose self-administration (**Fig 1A**). Alcohol and sucrose exposed mice responded an average of ∼100 times on the active lever (**Fig 1B**) reflecting an average preference ratio for the active lever of 81.3±5.8 and 86.1±2.8 percent, respectively (**Fig 1C**). Each group earned an overall average of 23 reinforcers per session (**Fig 1D**), which resulted in the alcohol group achieving an average intake of 0.83±0.12 g/kg/hr/day (**Fig 1E**). the sucrose group achieving an average intake of 11.4 mL/kg/day Overall, there were no behavioral differences between the two self-administration conditions.

**Figure 1.**
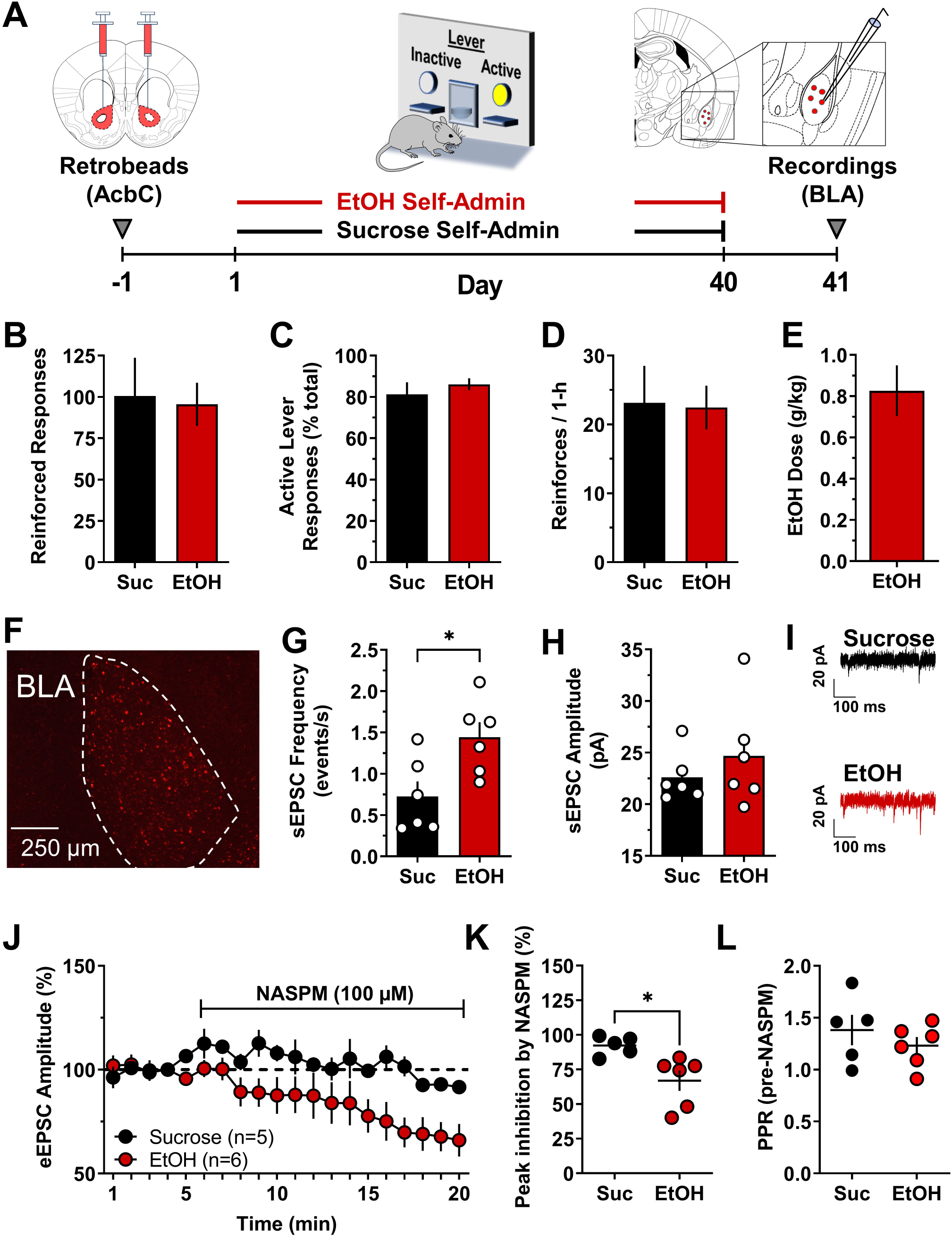
Enhanced CP-AMPAR activity in BLA neurons that project to the NAcb after alcohol self-administration as compared to sucrose. (**A**) Experimental timeline and schematic showing retrobead infusion in NAcbC followed by 40 days of ethanol (EtOH) or sucrose (Suc) self-administration. BLA recordings were conducted 24-h after the last self-administration session. (**B – E**) Parameters of EtOH (red bars) and sucrose (black bars) self-administration show behavior-matched performance between the sweetened alcohol and sucrose-only conditions with no differences in total responses (**B**), active lever responses (**C**), or reinforcers earned per hour (**D**), n=7/group. (**E**) Ethanol self-administration behavior resulted in average consumption of 0.83 g/kg per session. (**F**) Photomicrograph showing retrobead expression in the BLA following infusion in NAcbC. (**G**) Average plot showing a significant increase in sEPSC frequency in BLA neurons projecting to the NAcbC from alcohol exposed mice as compared to sucrose control; * - t(10) = 2.8, P = 0.02, alcohol n=6 cells from 6 mice, sucrose n=6 cells from 4 mice. (**H**) sEPSC amplitude was not changed by ethanol self-administration as compared to sucrose control. (**I**) Representative traces illustrating EtOH-induced increase in sEPSC frequency but not amplitude. (**J**) Time course of eEPSC amplitude (MEAN±SEM) during baseline (min 1 – 4) and bath application of NASPM (100 µM; min 5 - 20) from sucrose vs. ethanol self-administering mice. Data are normalized to minute 4 of the baseline. Alcohol n=6 cells from 6 mice, sucrose n=6 cells from 4 mice. (**K**) Peak inhibition of eEPSC amplitude by NASPM (MEAN±SEM from last 2 minutes of NASPM application) showing heightened sensitivity in EtOH self-administering as compared to sucrose control mice; * - t(9) = 2.5, P = 0.03. (**L**) MEAN±SEM paired-pulse ratio of eSPSC amplitude calculated during pre-NASPM baseline.

Retrobead infusion in the NAcb resulted in significant expression in the BLA, supporting the relevance of studying this direct pathway (**Fig 1F**). Recordings from retrobead-positive cells showed that alcohol self-administration was associated with a significant increase in sEPSC frequency with no change in amplitude as compared to sucrose controls (**Fig 1G-I**). The contribution of GluA2-lacking CP-AMPARs to synaptic transmission was determined by bath application of the CP-AMPAR blocker NASPM (100μM). NASPM produced a significant reduction in evoked EPSC (eEPSC) amplitude in retrobead-positive BLA neurons from alcohol self-administering, but not sucrose self-administering mice (**Fig 1J**). Aggregate data show that NASPM produced a peak 32% reduction in eEPSC amplitude in BLA→NAcb projection neurons from alcohol self-administering mice as compared to sucrose controls (**Fig 1K**), which is consistent with an alcohol-induced increase in post-synaptic expression of CP-AMPARs. To further assess the synaptic locus of alcohol-induced plasticity, we analyzed paired-pulse ratio (PPR) of eEPSCs, and the coefficient of variation (CV), prior to bath application of NASPM as indices of pre-synaptic glutamate release probability ^53,54^. Results showed no alcohol-induced change in mean PPR (**Fig 1L**) or CV (sucrose 1/CV=6.09±0.84; alcohol 1/CV=6.44±1.2). These data demonstrate that NAcb projecting BLA neurons express significantly more functional postsynaptic CP-AMPARs after 40 days of low-dose operant alcohol self-administration as compared to non-drug behavior-matched sucrose control.

### 3.2 CP-AMPAR activity in the BLA regulates operant alcohol self-administration

To assess potential CP-AMPAR regulation of the reinforcing effects of alcohol, we examined effects of site-specific infusion of the selective GluA2-lacking CP-AMPAR antagonist NASPM in the BLA prior to operant alcohol self-administration sessions (**Fig 2A)**. Bilateral infusion of NASPM dose-dependently (5.6 and 10 µg) decreased the positive reinforcing function of sweetened alcohol as measured by total responses on the alcohol-reinforced lever (**Fig 2B**). This finding is complimented by a dose dependent decrease in alcohol-reinforced response rate (**Fig 2C**) in the absence of effects on the inactive lever (not shown). NASPM also dose-dependently reduced alcohol dose (g/kg) consumed (**Fig 2D**). NASPM had no effect on the number of head pokes per reinforcer (**Fig 2E**), which indicates the absence of nonspecific changes in general consummatory behavior (e.g., ^29^). Mice emitted 57.3±20 (MEAN±SEM) total responses on the inactive lever (31% of total responding) under aCSF control conditions, and this was unchanged by NASPM. An effective dose of NASPM (5.6 µg) that reduced alcohol self-administration had no effect on total horizontal distance traveled (**Fig 2F**) or the velocity of activity (**Fig 2G**) in an open-field locomotor test suggesting that NASPM-induced reductions in alcohol self-administration were unrelated to nonspecific disruptions in motor activity or ability. Thus, these data show that NASPM infusion in the BLA reduced operant alcohol self-administration in the absence of nonspecific consummatory or motor effects, which suggests that the full expression of alcohol’s positive reinforcing function requires CP-AMPAR activity in the BLA.

**Figure 2.**
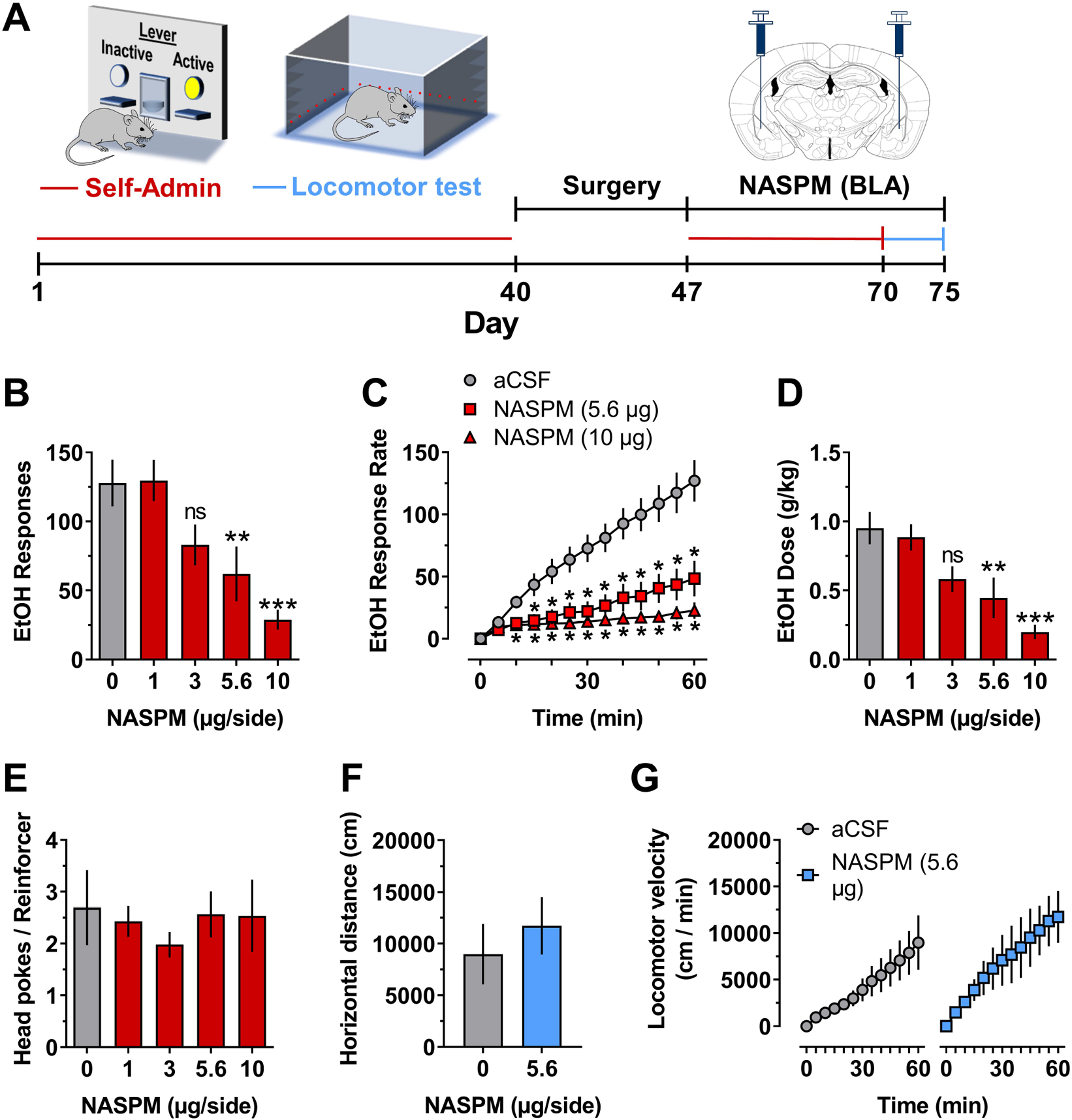
Blocking CP-AMPAR activity in the BLA reduced operant alcohol self-administration. (**A**) Timeline of procedures with schematic showing self-administration and locomotor test apparatus, duration of ethanol access, and anatomical location of NASPM infusions in the BLA. **B – E, Self-administration: (B)** NASPM (0 – 10 µg/side) infusion in the BLA significantly reduced total ethanol (EtOH) reinforced responses F(4,20) = 9.87, P < 0.0001, asterisks indicate significantly different from vehicle: ** - P = 0.006, *** - P < 0.0001; RM-ANOVA followed by Dunnett’s multiple comparison test. There was no individual (between subjects) difference in total responding F (5, 20) = 2.144, P = 0.1. (**C**) EtOH-reinforced response rate increased as a function of time F(12,60) = 32.3, P < 0.0001. NASPM (0, 5.6, and 10.0 µg/side) produced a significant dose-dependent reduction in EtOH-reinforced response rate: drug factor F(2,10) = 24.1, P = 0.0001, drug x time interaction F(24,120) = 26.5, P < 0.001, asterisks indicate significantly different from aCSF vehicle at corresponding time point: * - 0.002 ≤ P ≤ 0.04; RM-ANOVA followed by Dunnett’s multiple comparisons test. Response rate data were found to be normally distributed (D’Agostino-Pearson test, α=0.05) (**D**) EtOH dose (g/kg) consumed was reduced as a function of NASPM (0 – 10 µg/side) F(4,20) = 9.2, P = 0.0002, asterisks indicate significantly different from vehicle: ** - P = 0.01, *** - P < 0.001 Dunnett’s multiple comparison test. There were no between subject difference in EtOH dose F (5, 20) = 1.292, P=0.3064. (**E**) NASPM had no effect on number of head-pokes per reinforcer. **F – G, Locomotor activity:** NASPM (0 or 5.6 µg/side) infusion in the BLA had no effect on motor function as shown by total horizontal distance (cm/1-h) traveled (**F**) or velocity (cm/5-min) of locomotor activity (**G**). All data represent MEAN±SEM, n=6.

### 3.3 Alcohol increases GluA1-S831 phosphorylation in the BLA

GluA1-containing CP-AMPARs are activated (e.g., phosphorylated) at serine 831 (S831) in the amygdala by plasticity-inducing events ^55^. To evaluate the impact of alcohol self-administration on GluA1-containing AMPAR activation, parallel groups of C57BL/6J mice self-administered sweetened alcohol (alcohol 9% v/v + sucrose 2% w/v) or sucrose only (sucrose 2% w/v) during 30 1-h daily sessions (**Fig 3A**). Analysis of self-administration behavior showed no statistically significant group differences on measures of active lever presses (**Fig 3B**), percentage of responses on the active lever (**Fig 3C**), or number of reinforcers earned per hr (**Fig 3D**). Average (MEAN±SEM) alcohol intake across the 30 sessions was 0.92±0.11 g/kg/h (**Fig 3E**). Phosphorylation of GluA1-S831 was assessed in the BLA immediately after the last self-administration session. Operant alcohol self-administration was associated with a statistically significant increase in both the number of pGluA1-S831 immunopositive cells (**Fig 3F**) and pGluA1-S831 pixel density (**Fig 3G**) in the BLA as compared to the behavior-matched sucrose control group. Follow up qualitative analysis showed that GluA1 immunoreactivity was highly expressed throughout the BLA in both alcohol and sucrose exposed mice (**Fig 3H**) with expression in BLA neuronal membranes (**Fig 3I**), which is consistent with known expression of GluA1 during synaptic and behavioral plasticity ^56,57^. Thus, operant alcohol self-administration increases GluA1-S831 phosphorylation in the BLA of C57BL/6J mice to a greater extent that the nondrug reinforcer sucrose. This effect was observed under behavior-matched conditions indicating that differential GluA1-S831 phosphorylation was not related to differential operant conditioning-associated learning & memory, or motor activity during operant sessions.

**Figure 3.**
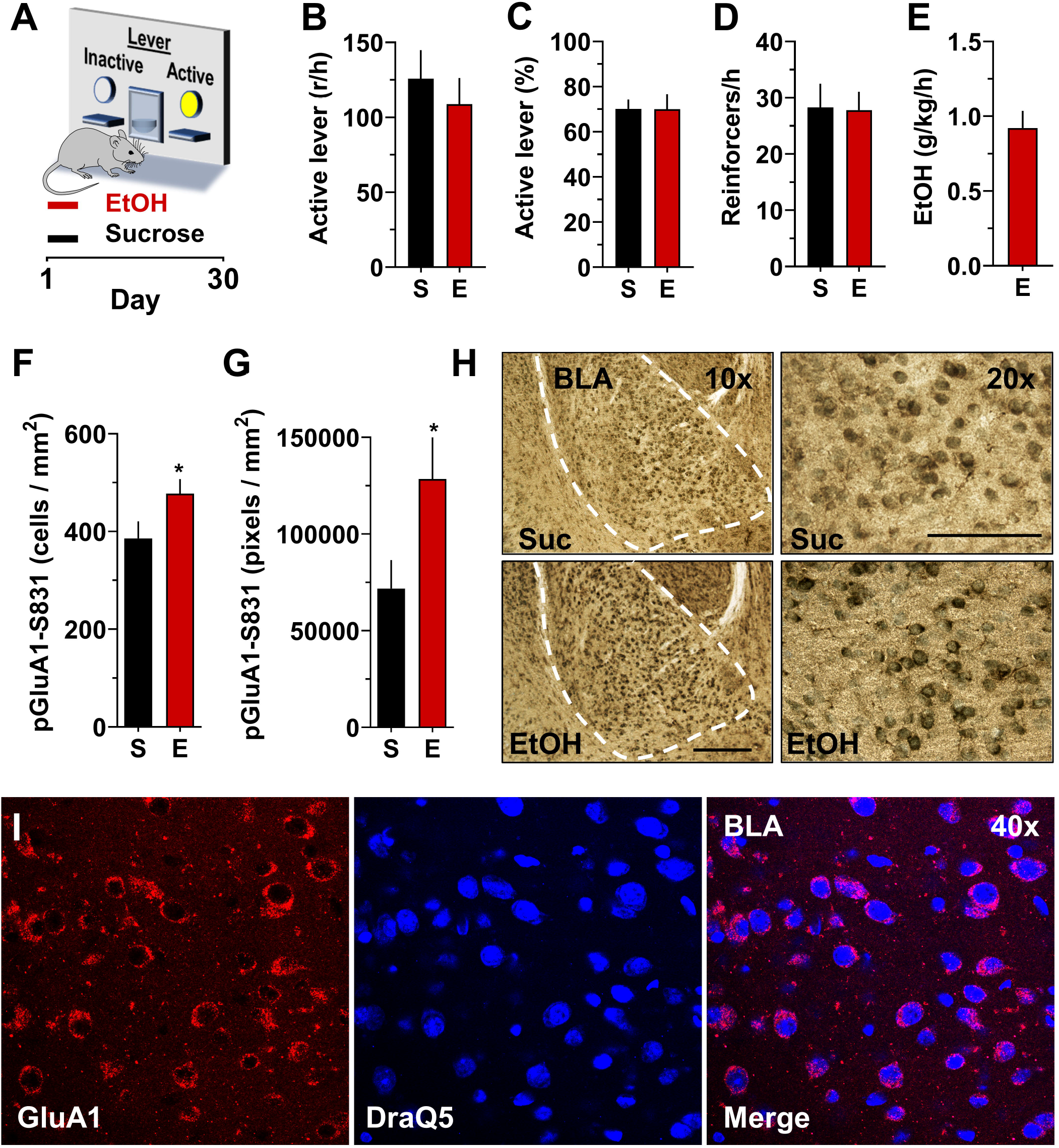
Operant alcohol self-administration increases pGluA1-S831 immunoreactivity in the BLA as compared to behavior-matched sucrose controls. **(A)** Schematic of operant conditioning chamber with active and inactive levers, and timeline of experimental procedure. (**B**) Number of responses per hour (r/h) on the active lever; (**C**) percentage of responses on the active lever showing preference; and (**D**) number of reinforcers earned per hour showed no difference between sucrose (S) and ethanol (E) self-administering mice. (**E**) Measurement of ethanol dosage (g/kg) showed that mice consumed pharmacologically meaningful levels of the drug during 1-h sessions as previously shown in Faccidomo et al., 2009. (**F-G**) Ethanol (E) self-administration was associated with a significant increase in pGluA1-S831 immunoreactivity (IR) as measured by cells/mm^2^ [t(14) = 2.0, P = 0.03] and pixels/mm^2^ [t(14) = 2.1, P = 0.026]. Data represent MEAN±SEM, n=8 / group; * - P<0.05 relative to sucrose (S) controls. (**H**) Photomicrographs showing pGluA1-S831 IR in the BLA after sucrose or ethanol self-administration at 10X (left) and 20X (right) magnification. Dashed line indicates the BLA boundary used for analysis of pGluA1-S831 immunoreactivity. (**I**) Photomicrographs showing illustrating the membrane-like nonoverlapping expression pattern of GluA1 compared to DraQ5 nuclear immunoreactivity in the BLA. All data represent MEAN±SEM, n=8.

### 3.4 GluA1ct expression in the BLA: Blocking AMPAR trafficking

Phosphorylation of GluA1 at S831 and other amino acid residues in the cytoplasmic c-terminus domain, including S845, plays a key role in activity-dependent synaptic trafficking of GluA1-containing AMPARs ^22,58,59^. To investigate the mechanistic role of AMPAR GluA1 trafficking in the positive reinforcing effects of ethanol, we infused a TET-ON viral vector in the BLA that encodes a dominant-negative GluA1 c-terminus (GluA1ct), which blocks activity-dependent synaptic delivery and function of GluA1-containing AMPARs ^36,37^ (**Fig 4A**). Doxycycline administration resulted in GluA1ct expression in the BLA with limited diffusion to adjacent nuclei (**Fig 4B**). GluA1ct expression was associated with significant reductions in the total number of alcohol-reinforced responses (**Fig 4C**) and alcohol intake (**Fig 4D**). We also evaluated alcohol reinforced response rate as a direct measure of positive reinforcement. Doxycycline administration significantly decreased alcohol-reinforced response rate in mice expressing GluA1ct in the BLA but was without effect in mCherry controls (**Fig 4E**). Significant decreases from baseline were observed after 10mg of Doxy starting at 20 – 60min. Importantly, alcohol-reinforced response rate was reduced after an initial response burst indicating normal onset of responding in the absence of nonspecific changes on memory or motor ability. MEAN±SEM responses on the inactive lever were unchanged by Doxycycline exposure (mCherry control = 20.94±4; GluA1ct = 20.96±7.5).

**Figure 4.**
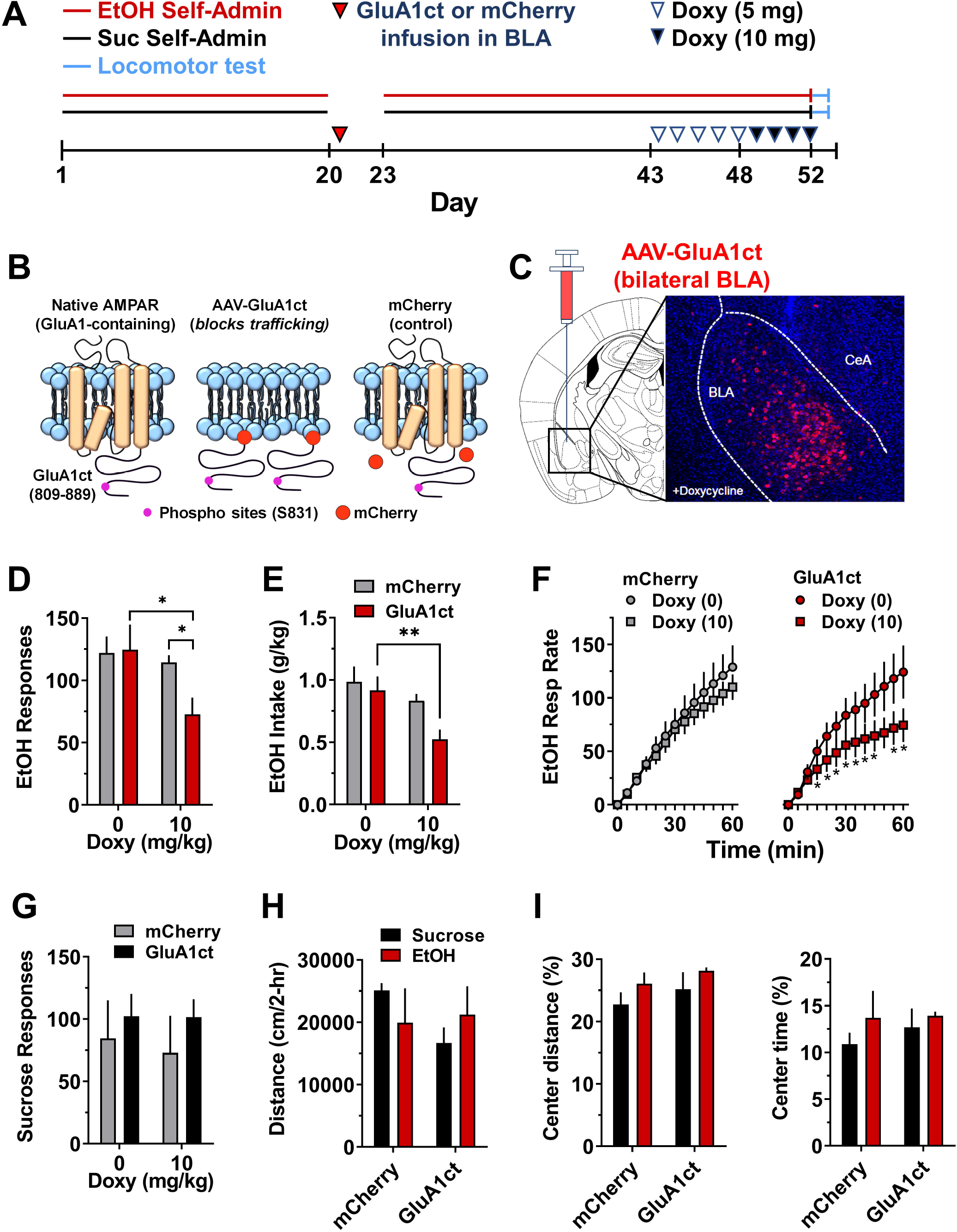
Inhibition of AMPAR GluA1 trafficking in the BLA selectively attenuated operant ethanol self-administration. (**A**) Timeline of experimental procedures. (**B**) Schematic showing native membrane bound GluA1-containing AMPAR with the c-terminus showing the CaMKII substrate S831, membrane block by the AAV-GluA1ct with mCherry tag, and mCherry control with normal AMPAR membrane trafficking. (**C**) Schematic showing region of the BLA with image of AAV-GluA1ct expression in the presence of doxycycline (Doxy). (**D**) Total ethanol-reinforced responses plotted as a function of oral Doxy dosage. Two-way RM-ANOVA – main effect of Doxy treatment: F(1,12) = 10.52; Doxy X GluA1ct interaction: F(1, 12) = 5.84; *P < 0.05 GluA1ct (n=6) versus mCherry control (n=8) after Doxy (10 mg/kg) and Doxy (0 mg/kg) versus Doxy (10 mg/kg) within GluA1ct; Šídák’s test. (**E**) Ethanol intake (g/kg) plotted as a function of Doxy. Two-way RM-ANOVA – main effect of Doxy: F(1, 12) = 13.6; P=0.003, ** - P < 0.01 planned t-test comparison of Doxy (0 vs. 10 mg/kg) within GluA1ct, Šídák’s test. (**F**) EtOH-reinforced response rate (cumulative resp / 5-min) plotted as a function of time (min). Separate two-way RM-ANOVAs – mCherry control group showed main effect of time: F (12, 168) = 83.5, P < .0001 but no effect of Doxy and no interaction. GluA1ct group showed main effect of time: F (12, 60) = 25.86, P<0.0001; main effect of Doxy: F (1, 5) = 8.139, P=0.0357; Doxy X time interaction: F (12, 60) = 5.68, P=0.044. Post hoc comparison: Doxy (10) versus Doxy (0) at corresponding time: *P<0.001 – 0.05. (**G**) Total sucrose-reinforced responses plotted as a function of Doxy dosage. RM-ANOVA found no effect of Doxy. (**H**) Distance traveled (cm/2-hr) in an open field by sucrose and ethanol self-administering mice plotted as a function of AAV condition (mCherry versus GluA1ct). (**I**) Thigmotaxis expressed as distance traveled in the center zone (% of total) and time spent in center (% of total) as a function of AAV condition. All data are presented as group Mean ± SEM.

To evaluate alcohol reinforcement-specificity, GluA1ct was overexpressed in BLA of behavior-matched sucrose controls. Doxycycline administration had no effect on the reinforcing effects of sucrose (**Fig 4F**), which suggests the absence of involvement of general reward or appetitive functions. Doxycycline produced no changes in MEAN±SEM responding on the inactive lever in sucrose controls (mCherry control = 16.1±1.5; GluA1ct = 16.8±6.7). Since the amygdala also regulates emotional responses that may impact alcohol intake, we assessed motor activity and thigmotaxis as an ethological measure of anxiety-like behavior in GluA1ct expressing alcohol and sucrose self-administering mice. Results showed no effect of GluA1ct expression on distance traveled (**Fig 4G**) or thigmotaxis (**Fig 4H**) in the open-field indicating normal motor activity and adaptive avoidance of open space in nondependent alcohol and sucrose self-administering mice. Thus, activity-dependent synaptic trafficking of GluA1-containing AMPARs in the BLA mediates the positive reinforcing effects of alcohol in a manner that is dissociated form general appetitive or emotional functions.

## 4 DISCUSSION

Exposure to alcohol and other drugs of abuse increases the expression and synaptic activity of GluA2-lacking CP-AMPARs in reward-related brain regions ^21,25^. However, the mechanistic role of CP-AMPARs in alcohol-seeking behavior has not been investigated. Results of this study show that sweetened alcohol self-administration increases expression of GluA2-lacking CP-AMPARs on BLA neurons that project to the NAcbC as compared to behavior-matched sucrose-only controls. Moreover, behavioral studies present compelling evidence that CP-AMPAR activity and GluA1 trafficking in the BLA mechanistically regulate operant alcohol self-administration. We propose that alcohol use promotes postsynaptic incorporation of CP-AMPARs in the BLA that, in turn, drive repetitive alcohol-seeking behavior during the development and progression of alcohol addiction.

The BLA sends excitatory glutamatergic projections to reward-related brain regions including the NAcb ^32^, suggesting that this structure relays information to the forebrain about reinforcing substances. Moreover, optogenetic activation of excitatory glutamate transmission in the BLA-to-NAcb neural circuit facilitates reward-seeking behavior in mice ^33^. To determine if operant alcohol self-administration increases CP-AMPAR synaptic activity of BLA neurons that project to the NAcb, we conducted whole-cell electrophysiology recordings from visually identified BLA cell bodies expressing retrobeads that were infused in the NAcb (e.g., BLA→NAcb projecting neurons) after sweetened alcohol versus sucrose self-administration. As lack of retrobead expression does not confirm that a particular BLA neuron does not project to the NAcb, we only recorded from retrobead positive neurons to identify the BLA→NAcb circuit. Importantly, alcohol and sucrose self-administration groups were behaviorally matched so that the impact of alcohol could be assessed in comparison to an equivalent history of non-drug reinforcement learning, memory, and motor activity associated with lever pressing. Alcohol self-administration was associated with an increase in sEPSC frequency with no change in amplitude, which could be consistent with an alcohol-induced increase in presynaptic glutamate release onto BLA→NAcb projection neurons ^60^; however, alterations in frequency of events could also be due to novel synapse synthesis due to the incorporation of CP-AMPARs into silent synapses ^61-63^. Because we fail to see a change in baseline CV or PPR between the groups, which would indicate presynaptic modulation, we believe the later interpretation is more consistent with our data.

To investigate potential incorporation of CP-AMPARS into synapses, we evaluated bath application of the selective GluA2-lacking AMPARs antagonist NASPM on retrobead labeled BLA neurons. NASPM decreased eEPSC amplitude in alcohol self-administering but not sucrose self-administering mice indicating that alcohol self-administration increases postsynaptic insertion of GluA2-lacking CP-AMPARs in BLA neurons that project to the NAcb in a manner that is unrelated to general nondrug reinforcement or motor function. This finding suggests that non-dependent alcohol self-administration is associated with enhanced plasticity at BLA-NAcb neurons ^64^ and may reflect an increase in the reinforcing properties of alcohol. Similar evidence shows that withdrawal from chronic intermittent ethanol ^25^ or cocaine ^19^ is associated with insertion of GluA2-lacking CP-AMPARs in the NAcb suggesting that upregulated CP-AMPAR expression can also occur in specific reward-related brain regions during multiple stages of addiction ^4^. By contrast, CP-AMPAR trafficking is not observed in paraventricular thalamus to NAcb synapses following cocaine withdrawal ^65^ and GluA1 surface expression is deceased in the NAcb following morphine exposure ^66^, indicating that upregulated CP-AMPAR trafficking is not a ubiquitous response to drugs of abuse. It is also important to note that glutamatergic synapses on interneurons express CP-AMPARs ^67^, indicating that NASPM application may alter synaptic activity of both BLA projection neurons and local inhibitory microcircuits, which may influence the reinforcing effects of alcohol ^51,68,69^.

Although these data demonstrate alcohol-induced changes in CP-AMPAR signaling in BLA-NAcb projection neurons, they do not address the generality of alcohol’s effects across multiple circuits. Topographically overlapping neurons in the BLA send glutamatergic projections to multiple regions that regulate aspects of reward and aversion, such as the insular cortex and central amygdala (CeA) ^70,71^, that may differentially alter alcohol self-administration ^72^. Future studies utilizing retrobead strategies can evaluate alterations in CP-AMPAR signaling in these and in other BLA projection regions. One interesting hypothesis is that CP-AMPAR activity would be downregulated by self-administered alcohol in BLA-CeA projecting neurons, which have been shown to undergo opposing synaptic changes as compared to BLA-NAcb projecting neurons following reward conditioning ^71^. A complementary approach would be to evaluate CP-AMPAR activity in a random population of retrobead-negative cells; however, differential synaptic changes across multiple unidentified circuits, and less than complete retrobead uptake, would complicate interpretation of results.

The finding that alcohol self-administration increases postsynaptic insertion of GluA2-lacking AMPARs in the BLA neurons raises the question as to whether this adaptation to alcohol has behavioral significance for drug-seeking behavior. Results show that site-specific infusion of the GluA2-lacking AMPAR antagonist NASPM in the BLA dose-dependently decreased alcohol reinforced response rate, which is a direct measure of reinforcement function. NASPM had no effect on general reward seeking (head pokes per reinforcer) or open field exploratory locomotor activity in a separate test. These pharmacological data show that the positive reinforcing effects of alcohol require GluA2-lacking CP-AMPAR activity in the BLA. Taken together with the electrophysiological data discussed above, these findings suggest that alcohol increases CP-AMPAR synaptic activity in the BLA and that this supports the positive reinforcing effects of the drug after prolonged non-dependent use. Thus, alcohol-induced upregulation of CP-AMPAR activity in the BLA may underlie the development and maintenance of repetitive drug use that characterizes the early stages of alcohol addiction.

CP-AMPAR regulation of synaptic plasticity is associated with activation (e.g., phosphorylation) of GluA1-S831 ^22,55^. To address the potential role of GluA1-containing CP-AMPAR activity in alcohol self-administration, we sought to determine if operant alcohol self-administration alters GluA1-S831 phosphorylation in the BLA. Non-dependent alcohol self-administration of sweetened alcohol produced a significant increase in pGluA1-S831 immunoreactivity in the BLA as compared to behavior-matched sucrose-only controls. This suggests that the increase in phosphorylation of GluA1-containing AMPARs is a pharmacological effect of alcohol, or related to specific rewarding effects of alcohol, rather than general reward processes or motor activity associated with lever pressing. This finding is consistent with evidence from a rat model of alcohol dependence showing that acute alcohol increases glutamate levels in the amygdala after chronic exposure to ethanol vapor ^73^. Elevated glutamate promotes Ca^2+^ influx, binding of Ca^2+^ by calmodulin, and rapid phosphorylation of CaMKII-T286 ^74^ which then phosphorylates AMPAR GluA1-S831, and other amino acid residues including S567, and promotes membrane insertion of the receptor during plasticity evoking events ^22,58,59^. Accordingly, GluA1-S831 and CaMKII-T286 phosphorylation are increased in the BLA of rats withdrawn from chronic ethanol vapor ^75^. Interestingly, elevated pGluA1-S831 was observed in the present study as an increase in both pixel density and number of immuno-positive cells, which suggests enhanced signaling both in active GluA1-containing cells and incorporation of previously inactive cells. We have shown that low-dose non-dependent operant alcohol self-administration increases both pCaMKII-T286 and pGluA1-S831 in the CeA, and that this effect is associated with upregulated AMPAR-mediated synaptic transmission ^14^. Together, these data suggest that alcohol targets GluA1-containing CP-AMPAR activity in the both the CeA and BLA amygdala during the development of alcohol addiction (e.g., self-administration) and expression of dependence (e.g., vapor exposure).

The subunit composition of AMPARs is a fundamental mechanism of activity-dependent plasticity ^76^. GluA2-lacking CP-AMPARs are rapidly recruited to synapses during plasticity ^22,23^. Thus, a compelling hypothesis is that GluA1 trafficking mediates maladaptive behavioral plasticity in addiction ^18,19,77-79^. To address this hypothesis, we tested the necessity of GluA1 trafficking for the positive reinforcing effects of alcohol by expressing an AAV that encodes a dominant-negative form of the GluA1 c-terminus that has been shown to prevent activity-dependent synaptic delivery of native GluA1-containing AMPARs via doxycycline-driven recombination ^36,37^. Results showed that doxycycline-dependent expression of the GluA1ct AAV in the BLA significantly decreased the positive reinforcing effects of alcohol as measured by total responding, response rate, and alcohol intake. There was no effect on sucrose self-administration in parallel controls, and no change in open-field locomotor activity or thigmotaxis, which is an index of anxiety-like behavior, suggesting that the impact of blocking GluA1-containing AMPAR trafficking was selective to alcohol self-administration. Thus, these data show for the first time that membrane trafficking of GluA1-containing CP-AMPARs in the BLA is required for the full expression of the positive reinforcing effects of alcohol.

It is noteworthy that although viral mediated expression of GluA1ct in the BLA reduced alcohol self-administration, the magnitude of effect was less than what was observed following infusion of the CP-AMPAR antagonist NASPM. This suggests that GluA1 trafficking was not fully inhibited by expression of dominant negative GluA1ct. However, prior evidence indicates that this AAV strategy virtually eliminates synaptic incorporation of GluA1-containing receptors in the lateral amygdala of naïve rats during LTP ^36^. Since we observed that alcohol self-administration during baseline both upregulated GluA1-S831 phosphorylation and promoted synaptic incorporation of CP-AMPARs in the BLA, it is plausible that a pool of functional postsynaptic CP-AMPARs remained in the membrane during doxycycline administration, which served to partially maintain behavior. This question could be addressed in a future line of investigation that evaluates the impact of alcohol self-administration on CP-AMPAR activity and synaptic incorporation in BLA neurons both before and after GluA1ct expression.

Importantly prior evidence indicates that global knockout of GluA1 does not alter alcohol drinking in mice ^80^; however, deletion of GluA1 in dopaminergic D1 cells prevents alcohol relapse-like behavior ^81^. This raises the possibility that mechanistic regulation of alcohol self-administration by GluA1ct expression seen in the present study may reflect a brain region-or cell-specific mechanism. Evaluating the role of GluA1 trafficking in other reward-related brain regions and specific cell types will be an important next step in understanding the generality of this mechanism of alcohol self-administration.

Expression of the dominant negative GluA1ct AAV in the BLA specifically reduced alcohol self-administration with no effect in parallel sucrose-only control mice. This is consistent with the NASPM-induced reduction in eEPSC amplitude only in alcohol self-administering mice, and the increase in pGluA1-S831 phosphorylation in the BLA from alcohol as compared to sucrose self-administering mice. Taken alone, these findings suggest that the reinforcing effects of sucrose are not mediated by CP-AMPARs in the BLA; however, other brain regions and specific genetic vulnerabilities may play a role in this process. For example, NASPM reduces AMPAR-mediated eEPSC amplitude in *ex vivo* NAcb slices from selectively bred obesity-prone, but not obesity-resistant, rats following exposure to a “junk food” diet ^82^. This suggests that CP-AMPAR expression in the NAcb may underlie “food addiction” that is associated with obesity ^83-85^. It would be interesting in future studies to evaluate the mechanistic role of CP-AMPAR trafficking in the NAcb in the regulation of sucrose self-administration to explore overlapping mechanisms of drug addiction and obesity.

In conclusion, results of the present study provide mechanistic evidence that CP-AMPAR activity (e.g., NASPM results) and GluA1 trafficking (e.g., GluA1ct AAV results) functionally regulate operant alcohol self-administration behavior. These data are consistent with a growing body of literature implicating CP-AMPAR activity in neural and behavioral pathologies associated with drug and alcohol addiction ^18-20,25,37,78,86^. Since the reinforcing effects of drugs contribute to the development and progression of addiction, CP-AMPAR activity in the BLA may be a critical mechanism that mediates pathology during initial and later stages of disease. It will be important to extend this line of investigation to other brain regions, such as the NAcb, to increase understanding of the neuroanatomical substrates of AMPAR-mediated behavioral pathologies in AUD. Moreover, it will be interesting to evaluate the potential mechanistic role of CP-AMPAR activity and GluA1 trafficking in relapse to alcohol-seeking behavior, which is regulated by AMPAR activity ^12,87,88^ and has significant translational value for clinical therapeutics ^89^.

## ACKNOWLEDGEMENTS

This research was supported by the National Institute on Alcohol Abuse and Alcoholism of the National Institutes of Health under award numbers R37 AA014983 (CWH) and P60AA011605 (CWH and ZAM) and by the Bowles Center for Alcohol Studies at The University of North Carolina at Chapel Hill. All authors report no financial interests or potential conflicts of interest. The authors wish to thank Dr. Manuel Mameli, University of Lucerne, Switzerland for kindly providing the GluA1ct AAV.

## AUTHOR CONTRIBUTIONS

CWH, ZAM, and SF designed the study. ZAM and ESC conducted electrophysiology experiments and analyzed the data. SF, JLH, OJH, MK, BLS, VRS, and SMT conducted behavioral, immunohistochemistry, western blot, viral vector, and site-specific pharmacological experiments and analyzed data. CWH, ZAM, SF, and JLH analyzed and coordinated final data presentation, and wrote the paper. All authors participated in interpreting the findings and approved the final version for publication.

## DATA ACCESSIBILITY

The database is available from the corresponding author upon request.

