## Supplemental Methods for "Calcium-Permeable AMPA Receptor Activity and GluA1 Trafficking in the Basolateral Amygdala Regulate the Positive Reinforcing Effects of Alcohol"

**Supplemental Materials and Methods**

**Animals**

Subjects were male C57BL/6J mice arriving at 10 weeks of age from Jackson Laboratory (Bar Harbor, ME). Mice were group-housed (n=4/cage) in clear, ventilated, techniplast cages (12” x 6.5” x 7”) lined with corn-cob bedding. A cylindrical PVC pipe and nestlet were provided for environmental enrichment. Purina rodent chow (Purina Isopro 3000) and water were available *ad libitum*, unless noted. The vivarium was maintained on a 12h:12h reverse light/dark cycle (lights off a 0800) with temperature at 21 ± 1 °C, and humidity at 40± 2%.

**Self-administration Apparatus and Procedure**

Self-administration sessions were conducted in computer controlled two-lever operant conditioning chambers (Med Associates, St. Albans, VT) as previously described ^1-5^. One week after arrival, mice were presented with alcohol (ethanol 9% v/v + sucrose 2% w/v) or sucrose only (sucrose 2% w/v) along with water, in their home cage for 2 weeks. We have found that initial home-cage access to the reinforcing solution familiarizes mice to the reinforcer and leads to reliable and consistent intake ^6-8^. All alcohol solutions (v/v) were prepared by diluting 95% ethanol (Pharmco Products Inc.; Brookfield, CT) with water.

After 2 weeks of home-cage intake, mice were fluid restricted for 23-hrs and placed in an operant conditioning chamber for a 16 h overnight session during which they performed an operant response (lever press) that was reinforced with a delivery of either the alcohol or sucrose-only solution into the drinking trough. A photobeam spanned the drinking trough which allowed for quantification of headpokes into the trough. We consistently find a positive correlation between the frequency of headpokes and reinforcements and the headpoke/reinforcer ratio can be used as a quantification of “drug seeking” behavior. The reinforcement volume was set at .014ml. During the first overnight session, mice responded on a fixed-ratio 1 (FR1) schedule of reinforcement. During a 2nd 16-hr session, the response requirement was increased from FR1 to FR2 to FR4. During a 3^rd^ overnight session, the response requirement was set at FR4 for the entire overnight session. Subsequent sessions were 1 h in duration and the response requirement was maintained at FR4. All 1 h sessions were conducted in the dark, between 1100–1600h, 6 days/week.

After every session, the drinking trough was checked to verify consumption of all delivered fluid. Our prior work has shown that the sweetened alcohol solution at a volume of 0.014 ml is fully consumed by free feeding C57BL/6J mice, which is required for valid demonstration of reinforcement function. Moreover, these procedures result in pharmacologically relevant blood alcohol concentrations of at the end of 1 h sessions (Salling et al., 2008; Faccidomo et al., 2009).

**Immunohistochemistry**

Following the 30th self-administration session, ethanol (n=8) and sucrose (n=8) self-administering mice were anesthetized and intracardially perfused with ice-cold phosphate buffered saline (PBS; 0.1M) followed by 4% paraformaldehyde. Coronal amygdala sections were then prepared for immunohistochemical analysis of pGluA1-Ser831 (1:200; PhosphoSolutions) as reported previously ^9-11^. A subset of tissue was prepared for immunofluorescent co-labeling of tGluA1-Ser831 (1:400; Abcam) and DraQ5 (1:2000; Abcam). Quantification of immunopositive cells and pixels in the BLA was performed using Bioquant Life Science Software (v. 8.40.20; Bioquant Corp., Nashville, TN) by an experimenter blind to treatment conditions. Pixel density and cell count measurements were calculated separately and divided by the area of the region and expressed as pixels/mm^2^ and cells/mm^2^, respectively. A confocal microscope (Leica TCS SL confocal microscope; Leica microsystems Inc.) was used to image and quantify the co-labeling of tGluA1-S831 and Draq5 at 40X.

**Stereotaxic Surgery**

Surgical procedures for retrobead injections, viral injections and cannulae implantation were conducted as follows. Mice were anesthetized with isoflurane (1-3% flow; 5 ml/kg) and placed into a stereotaxic frame (Kopf Instruments, Tujunga, CA). Mice were implanted with a 26-gauge injector guide cannula (Plastics One, Roanoke, VA) aimed bilaterally at either the BLA (NASPM and GluA1ct experiments) or N.AcbC (Nucleus Accumbens Core; retrobeads) using the following stereotaxic coordinates; BLA: -1.6 AP, ±3.0 ML, -3.9 DV from bregma; N.AcbC – (AP +1.6mm; ML ± 0.75mm; DV – 4.8mm; ^12^. Guide cannulae were positioned 2mm above the BLA and were secured to the skull with dental cement (Durelon, Butler Schein, Dublin, OH). A 33-gauge obturator (Plastics One) was inserted to prevent blockage. Mice were given ibuprofen (dose) for 3 days after surgery and were given 1 week to recover before starting or resuming behavioral experiments.

**Retrobeads & Patch-Clamp Electrophysiology**

All electrophysiology and analysis were performed blind to the treatment condition. In order to examine the BLA to N.AcbC projecting neurons in ethanol vs. sucrose self-administering mice, retrobeads (Red retrobeads IX, 10% (w/v) Lumafluor, Inc.) were directly injected (0.3 µl/injection; 0.1 µl/min flow rate) into ten-week old male C57BL.6J mice (n=16; ethanol (n=8), sucrose (n=8)). After recovery from surgery, mice were trained to perform an operant response for either sweetened alcohol or sucrose, as described above. Twenty-four hrs after the final 1 hr self-administration session (40-44 consecutive sessions/mouse), mice were anesthetized with isoflurane, rapidly decapitated and brains were prepared for slice physiology as previously described ^13,14^. Slices were prepared in a carbogenated (95% O2, 5% CO2) ice-cold sucrose artificial cerebral spinal fluid (sucrose-aCSF: 194 sucrose, 20 NaCl, 4.4 KCl, 2 CaCl_2_, 1 MgCl_2_, 1.2 NaH_2_PO_4_, 10 glucose, 26 NaHCO_3_). Following, slices were allowed to recover in carbogenated standard aCSF (ACSF: 124 mM NaCl, 4.4 mM KCl, 2 mM CaCl_2_, 1.2 mM MgSO_4_, 1 mM NaH_2_PO_4_, 10 mM glucose, 26 mM NaHCO_3_) at 32°C for at least 30 minutes, prior to transferring to the rig for recording. Slices were then transferred to a submersed recording chamber (Warner Instruments) and perfused with carbogenated aCSF (30°C) at 2 ml/min. Neurons in the BLA were visualized using infrared differential interference contrast (DIC) video-enhanced microscopy (Olympus). Whole-cell voltage-clamp recordings were conducted using borosilicate electrodes (3-5 MΩ) pulled with a P-97 Flaming-Brown micropipette puller (Sutter Instruments) filled with an internal solution. Signals were acquired via a Multiclamp 700B amplifier (Molecular Devices, Sunnyvale, CA, USA), digitized and analyzed via pClamp 10.6 software (Molecular Devices). Micropipettes were filled with a Cs-gluconate (Cs-gluc in mM: 135 Cs+-gluconate, 5 NaCl, 10 HEPES, 0.6 EGTA, 4 ATP, 0.4 GTP) internal solution for spontaneous EPSC (sEPSCs) recordings in the presence of picrotoxin (25 µM) to pharmacologically isolate AMPA receptor-mediated EPSCs. sEPSC recordings were acquired in 2-minute recording blocks at -80mV and were later analyzed using Mini Analysis (Synaptosoft). For evoked excitatory post-synaptic current (EPSCs) recordings, micropipettes were filled with a potassium gluconate (K-gluc) internal solution comprising (in mM): 70 K+-gluconate, 80 KCl, 1 EGTA, 5 HEPES, 2 MgATP, 0.3 GTP, and 1 Phosphocreatine and slices were perfused with normal, oxygenated aCSF. Evoked excitatory post-synaptic currents (eEPSCs) were evoked by local stimulation with Ni-chrome bipolar electrodes while neurons were voltage-clamped at −65 mV (the reversal potential for GABA-A, empirically determined in our preparation). This recording configuration was used to prevent oscillatory and polysynaptic events that are often observed in the BLA following conditioning (personal communication Roger Clem). After a 5-minute stable baseline was established, NASPM (100µM) was bath applied for 15 minutes while continuing to record eEPSCs, and recordings were later analyzed using Clampfit 10.6 software (Molecular Devices). Minutes 19-20 were used to compare drug treatments across groups.

To determine if alcohol-induced plasticity was modulated via pre-synaptic changes in glutamate release probability, the paired-pulse ratio (PPR) of eEPSPs recorded during baseline prior to bath application of NASPM was evaluated as previously described ^15^. Briefly, PPR was calculated as eEPSP2/eEPSP1, where eEPSP1 and eEPSP2 represent the amplitude of the first and the second eEPSP. Additionally, eEPSPs recorded during baseline prior to NASPM were used to derive the coefficient of variation (CV) as previously described ^15^. CV is a classical method for identifying the synaptic locus of changes in plasticity based on noise, or variability, due to transmitter release ^16,17^. CV was defined as as σ/μ where σ represents the mean amplitude and μ represents the standard deviation of 15 successive eEPSPs and plotted as 1/CV. CV can increase or decrease when plasticity is mediated by presynaptic mechanisms ^16,17^.

**NASPM Microinjections & Self-Administration**

To address whether BLA CP-AMPAR receptors regulate the reinforcing properties of alcohol, site-specific microinjections in the BLA of the selective CP-AMPAR antagonist NASPM (0 – 10 µg/side) were conducted in freely moving mice prior to self-administration sessions as reported previously ^6,11^. Briefly, male C57BL/6J mice (n=8) were trained to lever press for sweetened alcohol for 40 1-hr sessions. Next, they were implanted with a bilateral guide cannula into the BLA. Once stable levels of operant responding resumed after surgery, sham and then aCSF (vehicle; 0 dose) injections were conducted to habituate the mice to the handling required for site-specific microinjections. Two mice failed to reestablish baseline after surgery and were not used in the experiment leaving the sample size at n=6. Next, NASPM (0.5 μl/side; 0-10μg) was dissolved in aCSF and infused over a 4 min period (0.125μl/min) using a 1 μl Hamilton syringe connected to a Harvard Apparatus pump (Holliston, MA). A 33-gauge injector (Plastics One) extended 2mm beyond the tip of the guide cannula and remained in place for 1 min after the infusion ended to facilitate drug diffusion and to minimize vertical capillary action along the injection tract. Mice were unrestrained during the infusion and were placed immediately into the operant conditioning chamber for a 1 h session. Injections occurred maximally 2x/week. After completion of the dose-effect curve, mice were microinjected with either aCSF or 5.6 μg NASPM, in a counterbalanced design, and placed into an open field for a 1 hr session. The second locomotor test was conducted 7 days later. Immediately after the final microinjection session, mice were deeply anesthetized with sodium pentobarbital and were intracardially perfused with PBS and PFA for histological verification of injector placement in the BLA.

**GluA1ct Dominant-negative AAV & Self-Administration**

**Viral Construct**. In order to determine whether the positive reinforcing effects of alcohol require the phosphorylation of CP-AMPAR in the BLA, we used a Tet-On dominant-negative viral vector approach to competitively inhibit synaptic delivery of CP-AMPAR to the synaptic membrane. The adeno-associated virus (AAV) construct used in the present study (pAAV-TRE-GluA1ctmCherry-CMV-Tet-On3G; kindly provided by Manuel Mameli) was generated and authenticated as previously described ^18^. Briefly, the vector encodes a dominant-negative form of the GluA1 C-terminus (GluA1ct, corresponding to amino acids 809 to 889) that temporally prevents activity-dependent synaptic delivery of the AMPAR GluA1 subunit ^19^ via doxycycline-driven recombination ^20^. The AAV plasmids were made into recombinant AAV2/5 particles by the Vector Core Facility at the University of North Carolina, Chapel Hill, USA.

**Self-Administration and Microinjection of Viral Vector**. C57BL6J male (10-weeks old) were trained to perform an operant response for either sweetened alcohol or sucrose (n=16/group), and on Day 21 of baseline training, underwent stereotaxic surgery and were directly injected with either the GluA1ct AAV or an mCherry AAV (control group) into the BLA (see *Stereotaxic Surgery* for details). After 1 week of recovery, mice resumed operant self-administration training for 15 subsequent daily sessions, which allowed 22 days for AAV expression in the BLA. This expression period was chosen based on prior results with this AAV construct that showed significant AMPAR synaptic and behavioral effects follow a 21 day post-infusion period ^18^. The tet-on GluA1ct AAV was activated by administration of a palatable oral food pellet (POP) infused with doxycycline (0 or 10 mg/kg, b.i.d.). Doxycycline administration occurred times temporally distant from the self-administration session and consumption was verified by experimenter observation. All mice were first administered control (vehicle) POPs for 5 days followed by control or doxycycline for 10 subsequent days during which behavioral studies were conducted. After completion of the study, mice were deeply anesthetized, intracardially perfused, and brain were collected for histological verification.

**Locomotor Activity**

To determine if NASPM- or GluA1ct AAV-induced changes in operant behavior were associated with nonspecific motor effects, spontaneous locomotor activity was assessed as reported previously ^2,6,21-26^. Activity was measured in Plexiglas chambers (27.9 cm^2^; ENV-510, Med-Associates) via two sets of photobeams that recorded X–Y movements via computer interface. Ambulatory distance, velocity, and time spent in the center or periphery of the chambers were recorded at 100 ms intervals. Testing occurred for 60 min to be comparable to the timeline of the operant self-administration sessions.

**Statistical Analysis**

One-tailed t-tests were used to compare the behavioral outputs of operant ethanol and sucrose self-administering mice, differences in pGluA1 immunohistochemistry and dependent measures from electrophysiology experiments. For all electrophysiology experiments, investigators were blinded to the experimental group that the animals and cells were in until after all analysis was finished. For the NASPM dose-effect curves, one way RM ANOVAs were conducted for EtOH responses, dose & headpokes/reinforcer. Two-way RM Mixed Design ANOVAs were conducted to assess dose- and time- dependent effects on operant response rate and cumulative motor activity expressed as cumulative records (5-min bins). Behavioral significance for the AMPAR GluA1 trafficking experiment were assessed using two-way RM ANOVA s (AAV x DOXY). Post-hoc analyses were conducted when appropriate and the α-level was set at 0.05 for all statistical tests.
